# An ER retention motif controls the heteromeric stoichiometry of hERG1a/1b channels

**DOI:** 10.64898/2026.02.20.707017

**Authors:** Sudharsan Kannan, Liliana R. Ernandez, Gail A. Robertson

## Abstract

The *human ether-à-go-go related gene* (*hERG*) encodes a potassium channel essential for cardiac repolarization and neuronal excitability. In the heart, heteromeric assemblies of hERG1a and hERG1b subunits produce cardiac I_Kr_, and mutations in either subunit are associated with long QT syndrome. Although hERG1a and 1b contain identical transmembrane and C-terminal cytosolic domains, they differ in N-terminal cytosolic domains, with hERG1b harboring an arginine-based endoplasmic reticulum (ER) retention/retrieval motif that limits its surface expression in the absence of hERG1a. While the association of hERG1a and 1b subunits is known to influence channel function, the stoichiometry of heteromeric hERG channels and mechanisms that regulate it have remained unresolved. Here, using single molecule photobleaching step analysis in HeLa cells, we show that heteromeric hERG1a/1b channels assemble predominantly with a fixed 2:2 stoichiometry. Mutation of the hERG1b ER retention motif disrupts this bias, resulting in a broader, near-random distribution of subunit compositions. Independent functional assays using dominant-negative pore mutants in Xenopus oocytes yielded quantitative current suppression consistent with a 2:2 assembly and similarly revealed loss of stoichiometric bias upon RXR mutation. Together, these results establish the oligomeric composition of hERG1a/1b channels and identify ER retention as a previously unrecognized determinant of heteromeric stoichiometry.

**Statement of Significance:** Subunit stoichiometry is a key determinant of ion channel function, yet how defined stoichiometries are established during biogenesis remains poorly understood. Here we demonstrate that heteromeric hERG1a/1b channels assemble with a fixed 2:2 stoichiometry and that an arginine-based ER retention motif in hERG1b is required to maintain this assembly bias. Using single-molecule photobleaching and independent functional assays, we show that disrupting this motif leads to a broadened, near-random distribution of tetrameric subunit compositions. These findings reveal that ER retention contributes not only to quality control and trafficking but also specifies subunit stoichiometry during ion channel assembly.

## Introduction

The *human ether-à-go-go-related gene* (*hERG*, or *KCNH2*) encodes the Kv11.1 voltage-gated potassium channel, which is essential for repolarizing the ventricular action potential (1, 2). Loss of hERG function, through either genetic mutation or unintended channel block, prolongs the ventricular action potential and can lead to long-QT syndrome (LQTS) and sudden cardiac death (1–3). In addition, mutations and mis-splicing of *KCNH2* transcripts have been implicated in central nervous system disorders such as epilepsy and schizophrenia (4, 5). Thus, understanding the mechanisms that regulate hERG channel biogenesis is of broad biological and medical significance.

In human ventricular cardiomyocytes, alternate transcripts encode hERG1a and 1b subunits (ERG1a and 1b in other organisms), which co-assemble to form heterotetrameric channels that carry the repolarizing I_Kr_ current (6–11). The two subunits are identical except for their N-termini: ERG1a contains a Per-Arnt-Sim (PAS) domain (12), which is missing in ERG1b (6). Expressed independently in heterologous systems, ERG1a subunits form homomeric channels that efficiently traffic to the plasma membrane and produce potassium currents. In contrast, ERG1b homomers traffic poorly and produce little or no current (6, 13). Co-expression of ERG1b with ERG1a markedly alters channel gating, providing strong evidence for heteromeric assembly (6, 9, 14).

Multiple lines of evidence indicate that cardiac I_Kr_ is generated predominantly by heteromeric hERG1a/1b channels. The two subunits are expressed at approximately equal levels in canine and human myocardium and co-localize to T-tubules (8, 15). Computational and functional studies further show that hERG1b accelerates gating and promotes timely repolarization. In human induced pluripotent stem cell-derived cardiomyocytes (iPSC-CMs), loss of hERG1b – either by shRNA knockdown or by converting heteromeric channels to homomeric channels via trans application of a hERG1a N-terminal fragment (16–18)– prolongs and destabilizes action potential duration, producing a pro-arrhythmic phenotype consistent with computational predictions (7, 9). Consistent with this physiological role, mutations specific to hERG1b, like hERG1a, are associated with long QT syndrome and sudden cardiac death (9, 19).

Previous studies identified an N-terminal, arginine-based endoplasmic reticulum (ER) retention motif (RXR) in hERG1b that sequesters the subunit in the ER when expressed in the absence of hERG1a (13, 20). In heterologous expression systems, mutating the RXR motif effectively rescues hERG1b from ER retention: hERG1b proteins are fully glycosylated (mature) as assayed by western blot, and they produce measurable currents, indicating successful plasma membrane expression (13). This finding implies that ER retention plays a role in heteromerization of hERG1b and hERG1a. However, it is unknown whether ER retention merely permits heteromerization or actively determines the stoichiometry of hERG1a/1b channels.

Here, we used single-molecule photobleaching step analysis along with two-electrode voltage clamp in Xenopus oocytes to show that hERG1a/1b channels predominantly form 2:2 stoichiometry and that disrupting the hERG1b RXR ER retention motif eliminates this bias. Thus, ER retention does not simply permit assembly but actively defines subunit composition.

## Results

### Photobleaching step analysis confirms tetrameric stoichiometry of hERG1a homomers

To determine the subunit stoichiometry of hERG1a channels at the plasma membrane, we expressed GFP-tagged hERG1a in HeLa cells and performed total internal reflection fluorescence (TIRF) microscopy. To avoid signal contamination from overlapping hERG channels, we analyzed cells with moderate surface expression densities (<1 spot/μm²), minimizing the probability of two or more channels occupying a single diffraction-limited spot (∼250 nm) (Fig. S1A-B). We quantified 800-1000 discrete fluorescent puncta per condition and tracked their intensity decay over time to identify discrete photobleaching steps. Consistent with tetrameric architecture, cells expressing only hERG1a_GFP_ exhibited up to four photobleaching steps (Fig. 1. *B* and *C*).

**Fig. 1.**
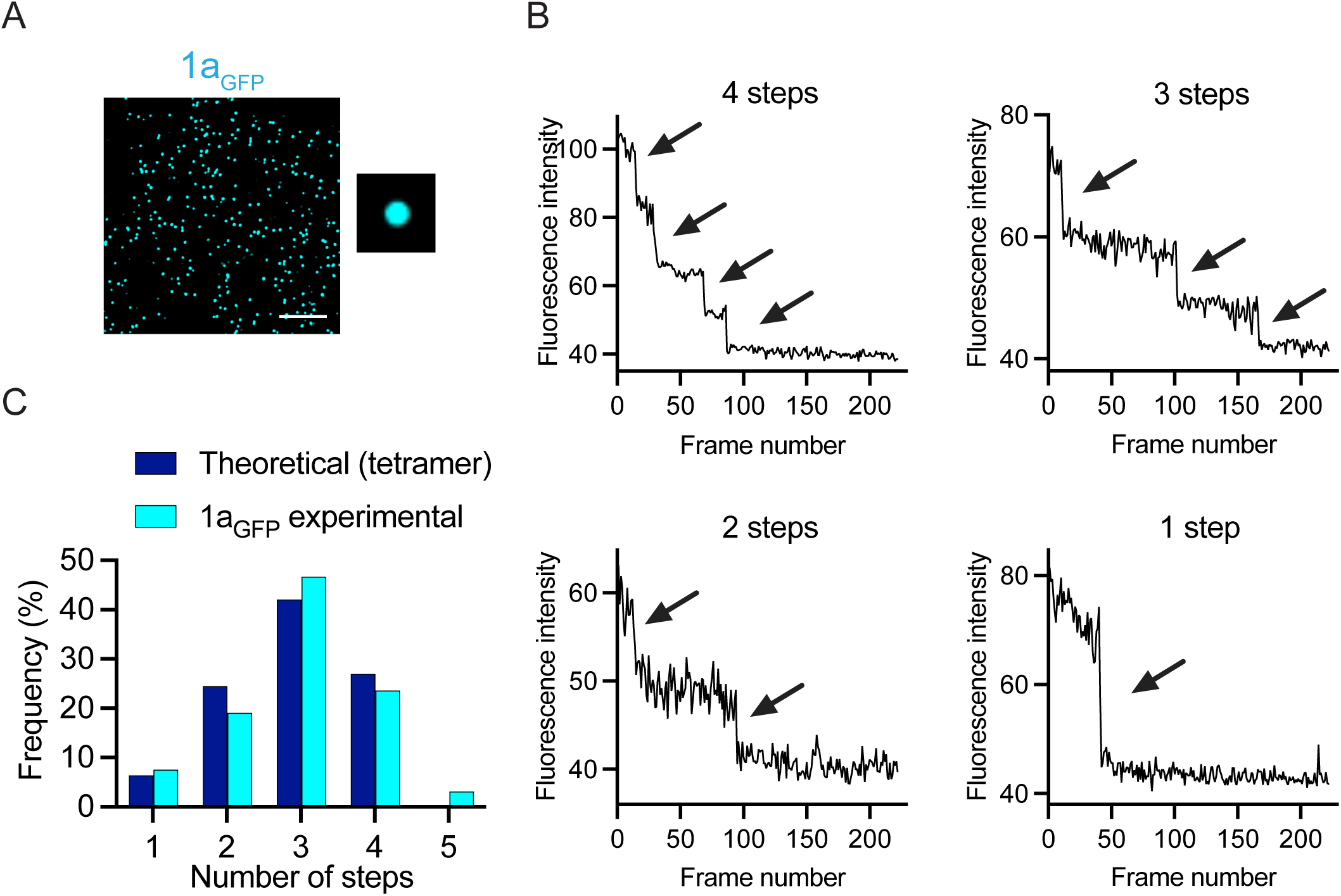
Photobleaching step analysis confirms tetrameric stoichiometry of hERG1a homomers at the plasma membrane. (*A*) TIRF image of HeLa cells transfected with plasmids expressing hERG1a_GFP;_ scale bar: 5 μm in large figure, inset frames is 1 μm width. (*B*) Sample fluorescent traces with four, three, two, and one photobleaching steps from cells expressing hERG1a_GFP_. (*C*) Photobleaching step distribution of hERG1a_GFP_ from ∼800 spots. The graph compares the experimentally observed distribution of 1-4 steps (cyan) with a theoretical step distribution corresponding to tetramers comprising 4 hERG1a_GFP_ subunits (blue) best fit with the theoretical distribution corresponding to 72% GFP detection efficiency.

Because the number of steps observed underestimates the true number of subunits per channel due to prebleaching and incomplete GFP maturation during biogenesis (21), we fit the observed photobleaching step distribution using a binomial model with varying GFP detection efficiencies. The best fit was achieved with a maturation probability of 72%, which was then used to generate a theoretical distribution that closely matched the experimental data (Fig. 1C). These results confirm that hERG1a channels form tetramers at the cell surface in agreement with previous structural studies (22, 23) and validate the use of photobleaching step analysis for estimating stoichiometry.

### hERG1a/1b heteromeric channels predominantly adopt 2:2 stoichiometry

To analyze the subunit stoichiometry of hERG1a/1b heterotetramers, we co-expressed 1a_GFP_ and 1b_SNAP_ in the same plasmid construct. We identified membrane-localized heteromeric channels by detecting GFP and SNAP-dye colocalized spots in TIRF images (Fig. 2A). Within these colocalized spots, recordings of GFP fluorescence as a function of time showed predominantly one or two photobleaching steps with fewer instances of three or four steps. The observed step distribution approximated the theoretical expectations for a tetramer containing two GFP-tagged subunits and was inconsistent with models representing other stoichiometries (Fig 2. B and C).

**Fig. 2.**
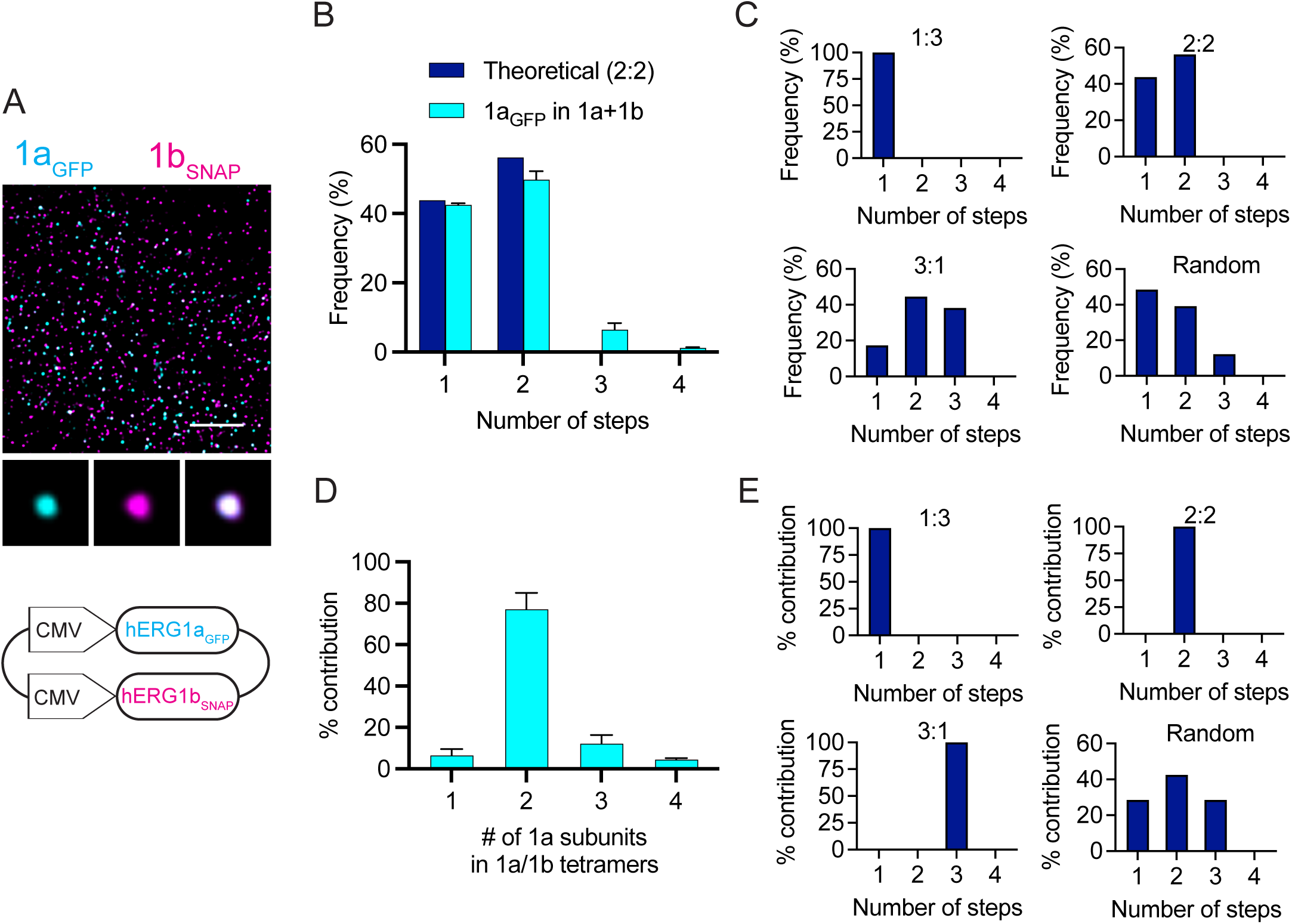
hERG1a/1b heteromeric channels predominantly form 2:2 stoichiometry. (A) TIRF image of HeLa cells transfected with single plasmid expressing hERG1a_GFP_ and hERG1b_SNAP_ together; scale bar: 5 μm in large figure, inset frames are 1 μm width. (*B*) Experimentally observed step distribution of 1a_GFP_ spots (blue) colocalized (centroids within 100 nm) with 1b_SNAP_ (cyan) showing close match with theoretical distributions for 2:2 (given 72% detection of GFP), analyzed from ∼800 spots. (*C*) Theoretical distributions for other possible stoichiometries for comparison to *B*. (*D*) the back-transformation of step distribution to oligomeric composition shows channels have a predominantly heterotetrameric stoichiometry of 2:2 hERG1a:1b. (*E*) Theoretical estimates of possible stoichiometry distributions for comparison to *D*.

Step distributions alone cannot unambiguously define stoichiometry, since similar patterns can arise from mixtures of channels containing different numbers of GFP-tagged hERG1a subunits. For example, a predominance of one- and two-step traces could reflect either a homogeneous population of assemblies comprising two hERG1a subunits or a heterogeneous mixture of assemblies containing one, two or three hERG1a subunits, particularly given incomplete GFP maturation. To distinguish between these possibilities, we modeled the observed step distribution as a weighted sum of binomial distributions corresponding to tetramers containing one, two, or three GFP-tagged subunits, explicitly accounting for the experimentally determined GFP detection efficiency (see Methods). Least-squares fitting revealed that the observed data are best explained by a dominant population of channels containing two hERG1a-GFP subunits (77%), with smaller fractions containing one (6%) or three (12%) subunits (Fig. 2D). This quantitative analysis demonstrates that hERG1a and 1b assemble predominantly in a 2:2 stoichiometry. Importantly, models of 1:3, 3:1, and random stoichiometry provided substantially poorer fits to the observed step distribution (Fig. 2E).

### Effects of the hERG1b ER retention motif on heteromeric stoichiometry

Having established that hERG1a/1b channels assemble predominantly with a fixed 2:2 stoichiometry, we next asked how this subunit composition is specified during biogenesis. Previous studies demonstrated that the hERG1b N terminus contains a canonical RXR-type ER retention motif that prevents forward trafficking of unpartnered hERG1b subunits, and a preferred association of hERG1b with hERG1a (13, 14, 20). Because heteromeric assembly occurs in the ER and involves N-terminal interactions between hERG1a and hERG1b, we reasoned that this retention motif might function not only as a trafficking checkpoint, but also as a determinant of heteromeric assembly stoichiometry.

To test whether disruption of the RXR motif alters heteromeric stoichiometry, we introduced the N^15^XN mutation into hERG1b and performed photobleaching step analysis of GFP-tagged hERG1a in co-expressing cells. Compared with wild-type channels, the photobleaching step distribution was markedly altered: the fraction of two-step events was reduced, with a corresponding increase in one-step and three-step events (Fig. 3C). Least-squares fitting of the step distributions revealed a substantial decrease in tetramers containing two hERG1a subunits and a concomitant increase in asymmetric 1:3 and 3:1 stoichiometries (Fig. 3D). Thus, disruption of the hERG1b ER retention motif compromises the strong bias toward 2:2 assembly observed in wild type heteromers, resulting in a broader distribution of subunit compositions.

**Fig. 3.**
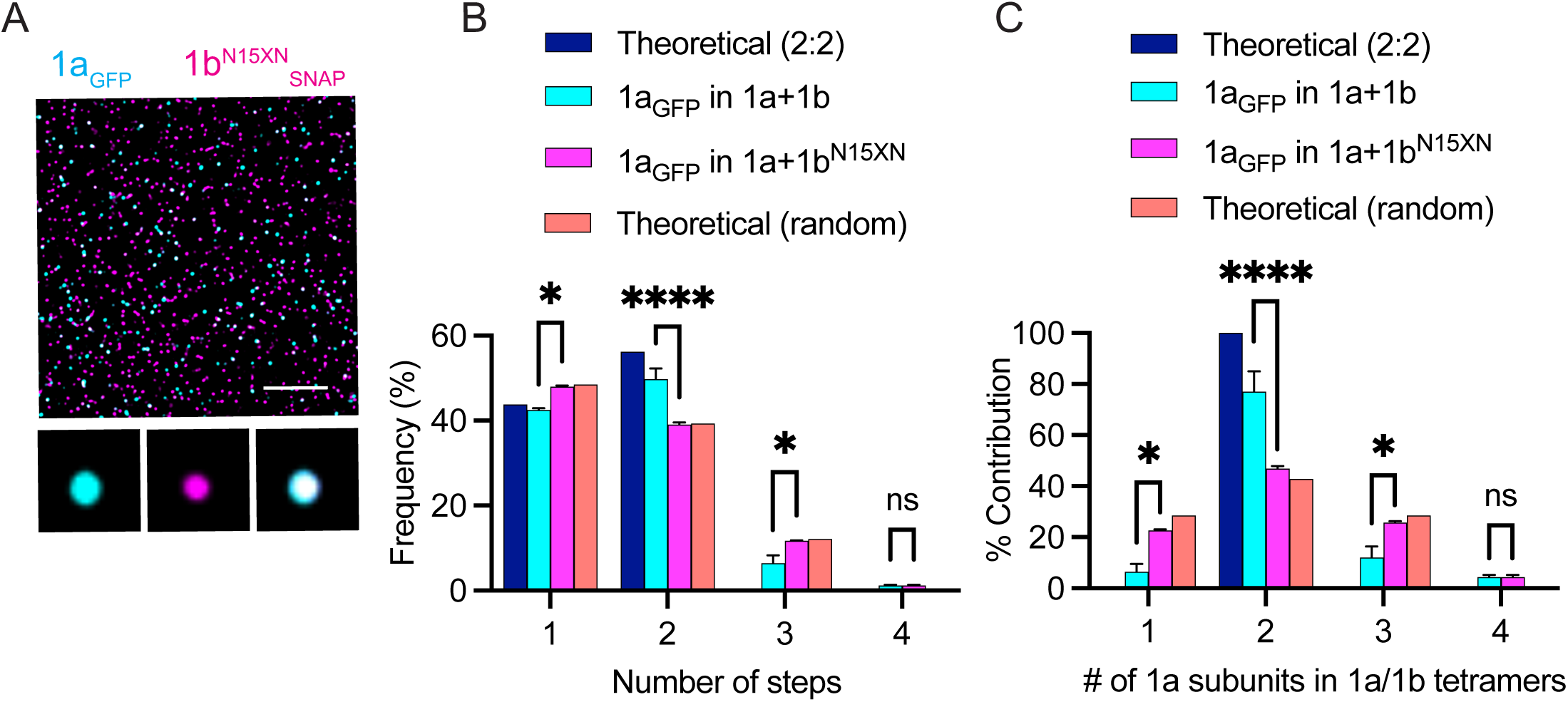
Mutating ER retention motif in hERG1b alters 1a:1b stoichiometry. (A) TIRF image of HeLa cells transfected with single plasmid expressing hERG1a_GFP_ and hERG1b_SNAP_^N15XN^ together. (*B*) Experimentally observed distribution of 1a_GFP_ (magenta) photobleaching steps in the spots colocalizing with 1b^N15XN^_SNAP_ compared with 1a expressed wild type 1b (cyan) and theoretical 2:2 (blue) and random (salmon). (*C*) The transformation of photobleaching steps to oligomeric composition shows that the NXN mutation data best fits with random distribution. For *C* and *D*, the results of Two-way ANOVA analysis showing differences in the photobleaching step and oligomeric distributions between 1a in 1a+1b and 1a+1b^N15XN^, p-values <0.0001 indicated by **** and <0.01 indicated by *. All photobleaching step-analysis shown in *C* and *D*are from a total of 800-1000 spots from n = 3 technical replicates (wells) for each condition. Data are mean ± S.E.M. Scale bar: 5 μm in large figures, inset frames are 1 μm width.

### Dominant-negative current suppression supports 2:2 hERG1a/1b stoichiometry

To determine hERG1a/1b stoichiometry using an independent functional assay, we exploited dominant-negative suppression by a non-conducting pore mutant, hERG1a-G628S (“poison”) (14, 24). Because hERG channels are tetramers, any assembled channel containing ≥1 poison subunit is expected to be nonfunctional. Thus, the fraction of remaining current after co-expression of WT and poison subunits depends on how subunits combine during assembly. Different assembly rules generate distinct quantitative relationships between injected RNA ratio and macroscopic current. We therefore calculated theoretical predictions for four assembly mechanisms: fixed 1:3, 2:2, or 3:1 stoichiometries, and random (binomial) stoichiometry (see Methods; Fig. 4A).

**Fig. 4.**
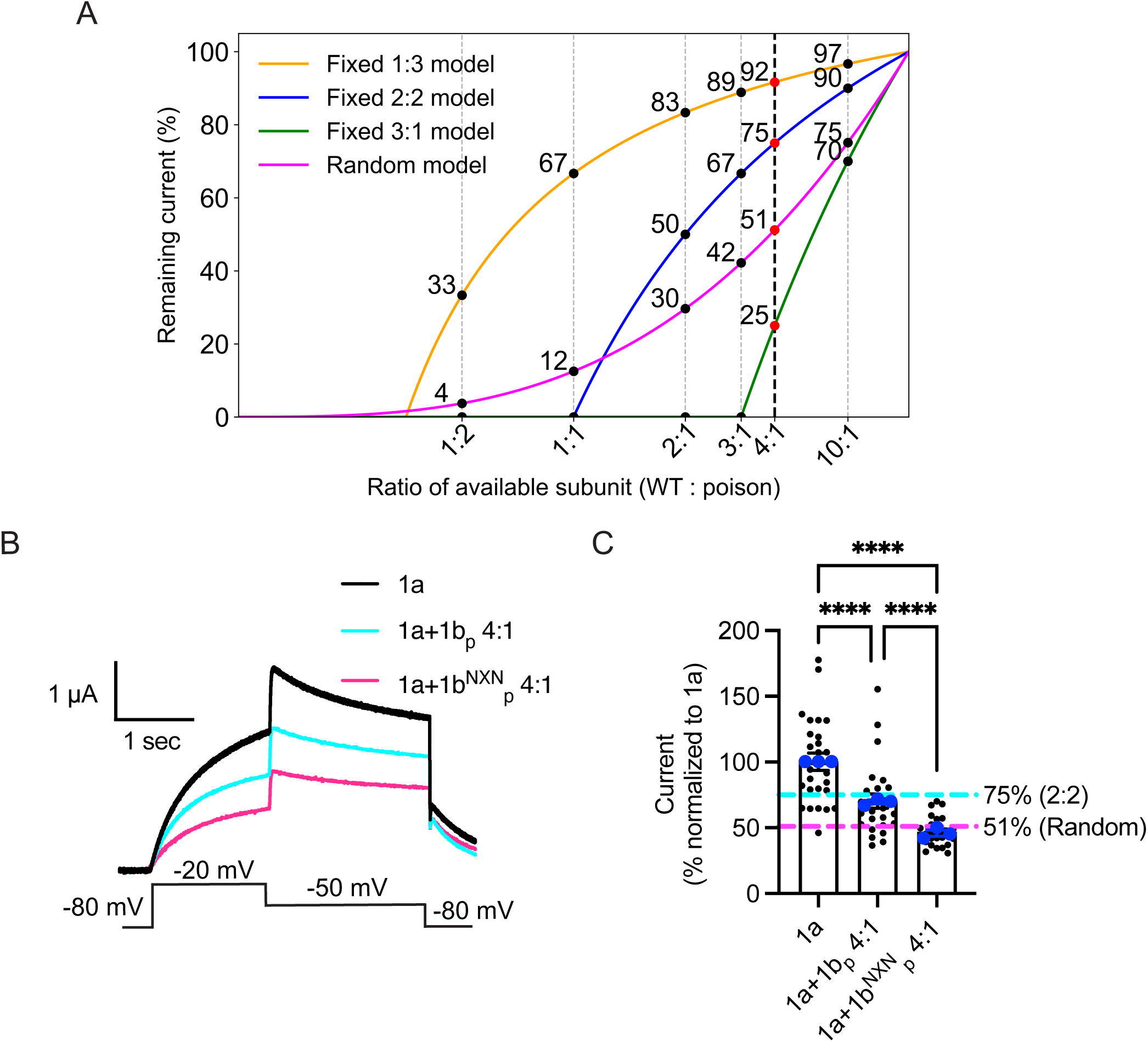
Dominant-negative current suppression confirms 2:2 hERG1a/1b stoichiometry. (A) Mathematical predictions of remaining current across varying WT:poison subunit availability ratios under various assembly models to predict the best RNA ratio that provides maximum separation. (B) Representative two-electrode voltage-clamp (TEVC) recordings from oocytes injected with hERG1a alone, or co-injected with hERG1b-poison or hERG1b^NXN^-poison at a 4:1 ratio (WT:poison). Voltage protocol and corresponding current traces are shown. (C) Peak outward currents were measured during the depolarizing step to −20 mV at the point of maximal current. Currents were normalized to the mean of the hERG1a-only condition within each experiment. Horizontal dashed lines indicate predicted current suppression assuming enforced 2:2 stoichiometry (75%, cyan) or random assembly (51%, magenta). Bars represent mean ± SEM from three independent replicates; blue points indicate replicate means, and black points indicate individual oocytes from all three replicates. Statistical comparisons were performed on replicate means using one-way ANOVA (**** p < 0.0001, ** p < 0.01, * p < 0.05, ns not significant).

To distinguish among assembly models, we injected Xenopus oocytes with a 4:1 ratio of hERG1a + hERG1b-poison subunits. This ratio was chosen because it most sensitively differentiates among models: random assembly predicts ∼51% remaining current compared to hERG1a currents, whereas the 2:2 fixed stoichiometry predicts ∼75% (Fig. 4A). Co-expression of hERG1b-poison with hERG1a reduced peak currents to ∼70%, approximating the 75% predicted for the 2:2 model (Fig. 4B-D). In contrast, co-expression of hERG1b-poison harboring the NXN mutation reduced currents to ∼47%, consistent with 51% predicted for random assembly (Fig. 4B-D). These results reinforce the conclusion that hERG1a/hERG1b heteromers preferentially assemble with a 2:2 stoichiometry, and that the NXN mutation disrupts this preference.

## Discussion

Subunit stoichiometry governs fundamental aspects of ion channel behavior, shaping surface expression, gating kinetics, and physiological output. Despite the central role of hERG channels in cardiac repolarization, the stoichiometry of hERG1a/1b heteromers had not previously been defined. Here, combining single-molecule photobleaching analysis with functional dominant-negative current suppression assays, we show that hERG1a/1b heteromers preferentially assemble with a 2:2 stoichiometry at the plasma membrane. Moreover, disruption of the hERG1b N-terminal ER retention motif abolishes this strong assembly bias, revealing a previously unappreciated link between channel biogenesis and mature channel stoichiometry.

While our results demonstrate that disruption of the hERG1b ER retention motif abolishes the bias toward 2:2 hERG1a/1b assembly, the mechanism by which this motif determines stoichiometry remains unclear. One possibility is kinetic control: wild-type subunits may be retained in the ER until proper assembly occurs, whereas disruption of the retention motif may allow premature ER exit and stochastic incorporation (13, 20). Alternatively, the NXN mutation may interfere with cotranslational association of hERG1a and 1b (25–27), shifting assembly toward post-translational interactions that yield random stoichiometry. A third possibility is that ER-resident chaperone proteins engage the hERG1b retention motif and actively guide subunit assembly toward a defined 1a:1b ratio. Consistent with this idea, SAP97 binds an arginine-rich ER retention signals in other receptors, and DNAJB proteins facilitate homotetrameric assembly of hERG1a; perhaps related processes participate in heteromeric hERG1a/1b assembly (28, 29).

Arginine-based ER retention motifs have long been recognized as critical quality-control elements in multimeric ion channels. Classic work on K_ATP_ channels demonstrated that an RKR motif present in both Kir6.x and SUR1 subunits restricts the trafficking of unassembled or partially assembled subunits, thereby ensuring only fully assembled, functional channels reach the plasma membrane (30). In the case of hERG1b, earlier studies established the presence of a canonical RXR motif and showed that is mutation permits surface expression of hERG1b homomeric channels (13). If the RXR motif functioned solely as a passive retention signal that became masked upon heteromerization, one would expect multiple heteromeric configurations, including both 2:2 and 3:1 configurations, to be represented at the cell surface. Instead, across independent experimental approaches, we consistently observe a strong bias toward a 2:2 hERG1a/1b stoichiometry. Disrupting the RXR motif abolishes this bias and permits a broader distribution of subunit compositions, revealing the RXR motif in hERG1b not only restricts incompletely assembled or homomeric channel surface expression but plays a previously unrecognized role in establishing a defined 2:2 stoichiometry in heteromeric hERG1a/1b channels.

Recent work from our group highlights additional complexity in hERG1a/1b biogenesis that informs the stoichiometric control described here. Using superresolution imaging, we showed that hERG1b expressed in isolation is sequestered within discrete, RXR-dependent ER subdomains that are distinct from degradative quality-control pathways (20). Introduction of hERG1a into cells containing these preformed hERG1b puncta leads to their dissolution and rescue of previously synthesized hERG1b, indicating that a pool of assembly-competent hERG1b can be preserved in the ER. How this sequestration pathway relates to cotranslational heteromerization remains unresolved. One possibility is that RXR-dependent sequestration functions as a buffering mechanism when hERG1b synthesis exceeds the availability of hERG1a, capturing excess subunits that are not cotranslationally incorporated. Interestingly, the rescued subunits associate posttranslationally, rather than cotranslationally, but the stoichiometry of the rescued channels remains to be determined. In this framework, stoichiometric control may reflect not only subunit recognition, but also the kinetics of subunit supply and ER handling.

A recurring concern in single-molecule photobleaching studies is that fluorescent protein tagging or plasmid-driven co-expression could bias subunit incorporation and artificially constrain apparent stoichiometry. Several observations argue against this interpretation in the present study. First, mutation of the hERG1b ER retention motif markedly alters the photobleaching step distribution, despite identical tagging and expression strategies, indicating that neither fluorescent labeling nor plasmid design alone dictates the observed 2:2 stoichiometry. Second, independent functional assays using untagged subunits yield quantitatively consistent results. Under a 4:1 hERG1a: hERG1b expression ratio, dominant-negative suppression produces current levels closely matching predictions for 2:2 channel assembly and inconsistent with expectations from purely expression-driven or random incorporation models. Together, the concordance between single-molecule imaging and functional measurements supports the conclusion that 2:2 stoichiometry reflects intrinsic features of hERG1a/1b assembly rather than experimental artifact.

Together, these findings establish that heteromeric hERG1a/1b channels assemble with a defined 2:2 stoichiometry and that this assembly bias depends on an arginine-based ER retention motif within the hERG1b N terminus. By showing that disruption of this motif leads to loss of stoichiometric constraint without abolishing heteromerization, our work reveals subunit stoichiometry as a regulated outcome of channel biogenesis rather than a passive consequence of expression levels. Given evidence from human iPSC-derived cardiomyocytes and computational models that loss of hERG1b destabilizes repolarization and promotes arrhythmogenic behavior, defects in stoichiometric control may represent an underappreciated mechanism contributing to cardiac disease. More broadly, these results suggest that ER retention motifs can shape not only channel trafficking but also the composition of multimeric membrane proteins, adding a new dimension to the regulation of excitability in health and disease.

## Methods

### Cell culture

HeLa cells (ATCC, CCL-2) were cultured in 35mm glass bottom dishes with #1.5 cover glass (Cellvis, D35-14-1.5-N) using DMEM culture media (ThermoFisher, 11995065) supplemented with 10% FBS (ATCC, SCRR-30-2020). HeLa cells were transfected with 400 ng of plasmid per 35mm dish with 2 ul of TransIT-X2 Transfection reagent (Mirus Bio, MIR6004), and cells were imaged after 24 h of transfection. 100 nM SNAP-tag ligand JF585 was added to the culture media for 30 min, followed by three washes for 5 min each using pre-warmed culture media to remove the free ligand.

### Plasmid constructs

Overexpression plasmids were made from pcDNA3.1 as a backbone. Truncations and insertion of coding sequences of hERG and fluorescent proteins were performed using In-Fusion® Snap Assembly Master Mix (Takara, 638948). For heteromeric stoichiometry experiments, we expressed hERG1a_GFP_ and hERG1b_SNAP_ under two distinct CMV promoters in the opposite direction to express relatively the same level of hERG1a and 1b.

### Microscopy

For single-molecule photobleaching analysis, cells that sparsely expressed hERG protein were imaged with a Nikon Ti2 microscope equipped with TIRF 100x (1.45NA) oil immersion objective, a Hamamatsu ORCA-Flash4.0 sCMOS camera, and an iLas2 total internal reflection fluorescence (TIRF) imaging system. Sequential imaging of GFP and SNAP-tagged subunits was performed using 488 nm and 561 nm laser excitation, respectively. Images were acquired continuously at 50 ms per frame for a total of 2 minutes without inter-frame delay.

### Estimating stoichiometry from fluorescent data

Cells exhibiting moderate surface expression of hERG (<1 spot per µm^2^) were selected to minimize the likelihood of multiple fluorescent subunits occupying a single diffraction-limited spot (see Fig. S1).

Fluorescent puncta detected automatically with Imaris software (Oxford Instruments plc) were analyzed only if they showed no evidence of blinking or spatial overlap with neighboring spots. Spots exhibiting multiple Gaussian peaks and signs of fluorophore clustering were excluded from analysis. The first four frames of images were merged and deconvolved using Huygens Professional software (Scientific Volume Imaging), using the Classic Maximum Likelihood Estimation (CMLE) algorithm with default settings. Deconvolved images were segmented using the Imaris spots model, and masks were exported. The masks were applied to the raw images the intensity was measured across time in Fiji (ImageJ). The fluorescence intensity was fitted by recursive partitioning using the ‘rpart’ package in R to identify the number of photobleaching steps in each fluorescent trace in an automated manner. Given that the GFP detection efficiency is not 100%, the GFP detection rate in our experimental condition, i.e., parameter *m* of the binomial distribution, was estimated by the least-squared estimation. This involves a) calculating theoretical probabilities of various outcomes based on different probability values, b) comparing this with the observed photobleaching steps, and c) identifying the probability values that resulted in the smallest discrepancy between the theoretical and observed data. This approach allows us to estimate the most accurate GFP detection probability (*m*), which we found to be 0.72, similar to that reported in previous studies utilizing this approach (21).

To predict which combination of 1a subunits (1, 2, or 3) in hERG1a/1b heterotetramers would result in the observed photobleaching step distribution, we used the least squares approach modified from previously published method (21). To estimate the best-matched percentages of the step distribution that fit the observed fractions of steps, we generated the theoretical binomial distributions for each combination of the number of subunits (*n* with *n_max_*= 4) and the number of photobleaching steps (*k*) with a success probability (*m*) of 0.72, accounting for the observed GFP detection efficiency. The number of subunits n was taken from 1, as the zero photobleaching steps cannot be experimentally observed. We used the binomial probability formula:

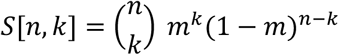

S would result in a matrix with each row representing a different number of subunits (*n*) and each column representing the number of photobleaching events (*k*):

| $S [k, n]$ | $n= 1$ | $n= 2$ | $n= 3$ | $n= 4$ |
| --- | --- | --- | --- | --- |
| $k= 1$ | 0.72 | 0.4032 | 0.1693 | 0.0632 |
| $k= 2$ | 0 | 0.5184 | 0.4354 | 0.2438 |
| $k= 3$ | 0 | 0 | 0.3732 | 0.4180 |
| $k= 4$ | 0 | 0 | 0 | 0.2687 |

As we did not count zero photobleaching steps (*k*= 0), we normalized the matrix *S* by summing up each row of *S* to obtain the normalized matrix:

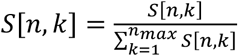

We then formulated a system of linear equations:

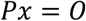

where 0 is the transposed matrix of theoretical distributions (!), 2 is the vector of observed fractions of steps, and 1 is the vector representing the fraction of each subunit configuration. We solved for 1 using the least square (normal equation) formula below, which provides the final estimates for the fractions of each subunit configuration:

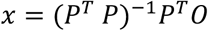

### Voltage clamp recordings

pGH19 plasmids containing hERG CDS with T7 promoter were linearized using NotI-HF restriction enzyme (NEB, R3189L) at the 3’ end. Linearized plasmids were transcribed using mMESSAGE mMACHINE™ T7 Transcription Kit (Invitrogen, AM1344) and purified using MEGAclear™ Transcription Clean-Up Kit (Invitrogen, AM1908) and 10ng cRNA was injected into oocytes. For poison-subunit assays, the non-conducting pore mutant hERG1a-G628S (hERG1a-poison) and the equivalent hERG1b (G288S) constructs (hERG1b-poison or hERG1b-NXN-poison) were co-injected with wild-type hERG1a at a fixed ratio of 4:1 (WT: poison). Oocytes were maintained at 18 °C in ND96 solution and used for recordings 1-2 days after injection.

Whole-cell currents were recorded using two-electrode voltage clamp (TEVC) at room temperature with electrodes filled with 2 M KCl. Currents were elicited by depolarizing steps from −80 mV to -20 mV (2 s duration) followed by repolarization to −50 mV. Peak outward currents were measured at the end of the depolarizing step to −20 mV, a voltage that has been shown to elicit maximal outward hERG1a currents while minimizing residual currents from non-conducting poison subunits (14).

### Theoretical predictions for poison-subunit suppression

We modeled tetrameric assembly under the assumption that functional current requires incorporation of four wildtype (WT) subunits, and that any tetramer containing one or more poison subunits is non-conducting. Let f represent the fraction of WT subunits in the total assembly pool:

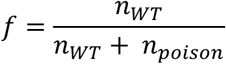

where *n*_*WT*_ is number of wild-type subunit and *n*_*poison*_ is number of poison subunits.

In the following assembly models, each model assumes heteromers are non-conducting and only leftover WT subunits can form conducting tetramers. The number of leftover WT subunits is *n*_*WT*,*left*_ and they can form up to 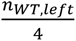 conducting tetramers. To compare the conducting tetramers with the WT-only condition, we normalize the current as:

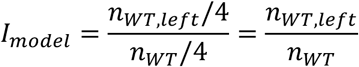

#### A. Fixed 1:3 assembly

In the 1:3 model, each heterotetramer is constrained to comprise exactly 1 wildtype (WT) subunits and 3 poison subunits when both are available. The number of leftover WT subunits is 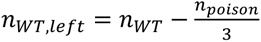.

The normalized current for 1:3 model is,

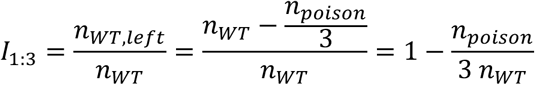

Expressed in terms of the WT fraction *f* (defined above), this becomes:

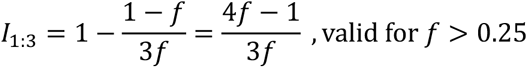

For 4:1 WT:poison ratio (*f* = 0.8), this predicts:

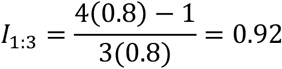

#### B. Fixed 2:2 assembly

In the 2:2 model, each heterotetramer is constrained to comprise exactly 2 wildtype (WT) subunits and 2 poison subunits when both are available. The number of leftover WT subunits is *n*_*WT*,*left*_ = *n*_*WT*_ − *n*_*poison*_.

The normalized current for 2:2 model is,

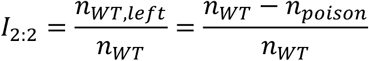

Expressed in terms of the WT fraction *f* (defined above), this becomes:

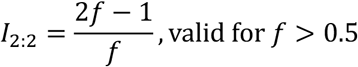

For 4:1 WT:poison ratio (*f* = 0.8), this predicts:

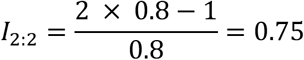

Thus, 75% of the WT current is predicted to remain under enforced 2:2 assembly.

#### C. Fixed 3:1 assembly

In the 3:1 model, each heterotetramer is constrained to comprise exactly 3 wildtype (WT) subunits and 1 poison subunits when both are available. The number of leftover WT subunits is *n*_*WT*,*left*_ = *n*_*WT*_ − 3 *n*_*poison*_. The normalized current for 3:1 model is,

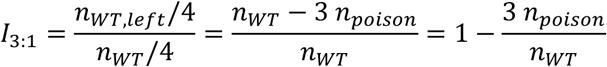

Expressed in terms of the WT fraction *f* (defined above), this becomes:

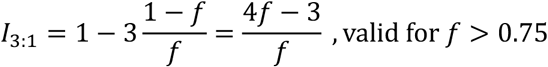

For 4:1 WT:poison ratio (*f* = 0.8), this predicts:

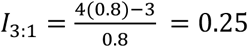

#### D. Random (binomial) assembly

If tetramers assemble by independently selecting four subunits from a large pool (approximating sampling with replacement), the probability of exactly *k* poison subunits follows the binomial distribution:

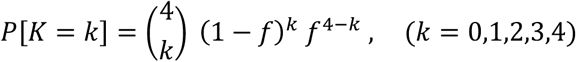

Only tetramers with no poison subunits (i.e., k = 0) are conducting. Thus, the probability that a tetramer is functional is:

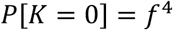

To compare the current in a poison condition to the WT-only baseline using the same number of WT subunits, we compute the relative current:

1. WT-only: total number of conducting tetramers is *n*_*WT*_/4
2. Mixed (poison): expected number of conducting tetramers is 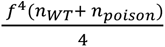.

Taking their ratio yields:

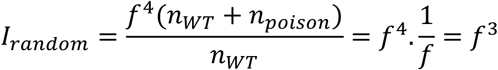

Thus, the predicted current (normalized to WT-only) under random assembly is:

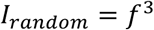

For a 4:1 injection ratio of WT:poison subunits:

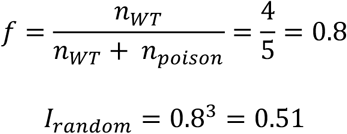

Therefore, random assembly predicts ∼51% of WT current would remain in the presence of poison subunits at this ratio.

| Model | Fixed 1:3 | Fixed 2:2 | Fixed 3:1 | Random (Binomial) |
| --- | --- | --- | --- | --- |
| Predicted I ($f = 0.8$) | 0.92 | 0.75 | 0.25 | 0.51 |

## Acknowledgments

We thank members of the Robertson lab, especially Drs. Catherine Eichel, Lisandra Flores-Aldama, and Taylor Voelker for critical discussion, Fang Liu for technical support, and Dr. Anjon Audhya and members of his lab for use of equipment and helpful advice. We thank Dr. Luke D. Lavis of Janelia Farms for providing Janelia Fluor dyes. The work was supported by NIH grants R01-HL131403 and R01-NS081320 (G.A.R.) and American Heart Association Predoctoral Fellowship 24PRE1188958 (S.K.).

## Supplementary Information

**Fig. S1.**
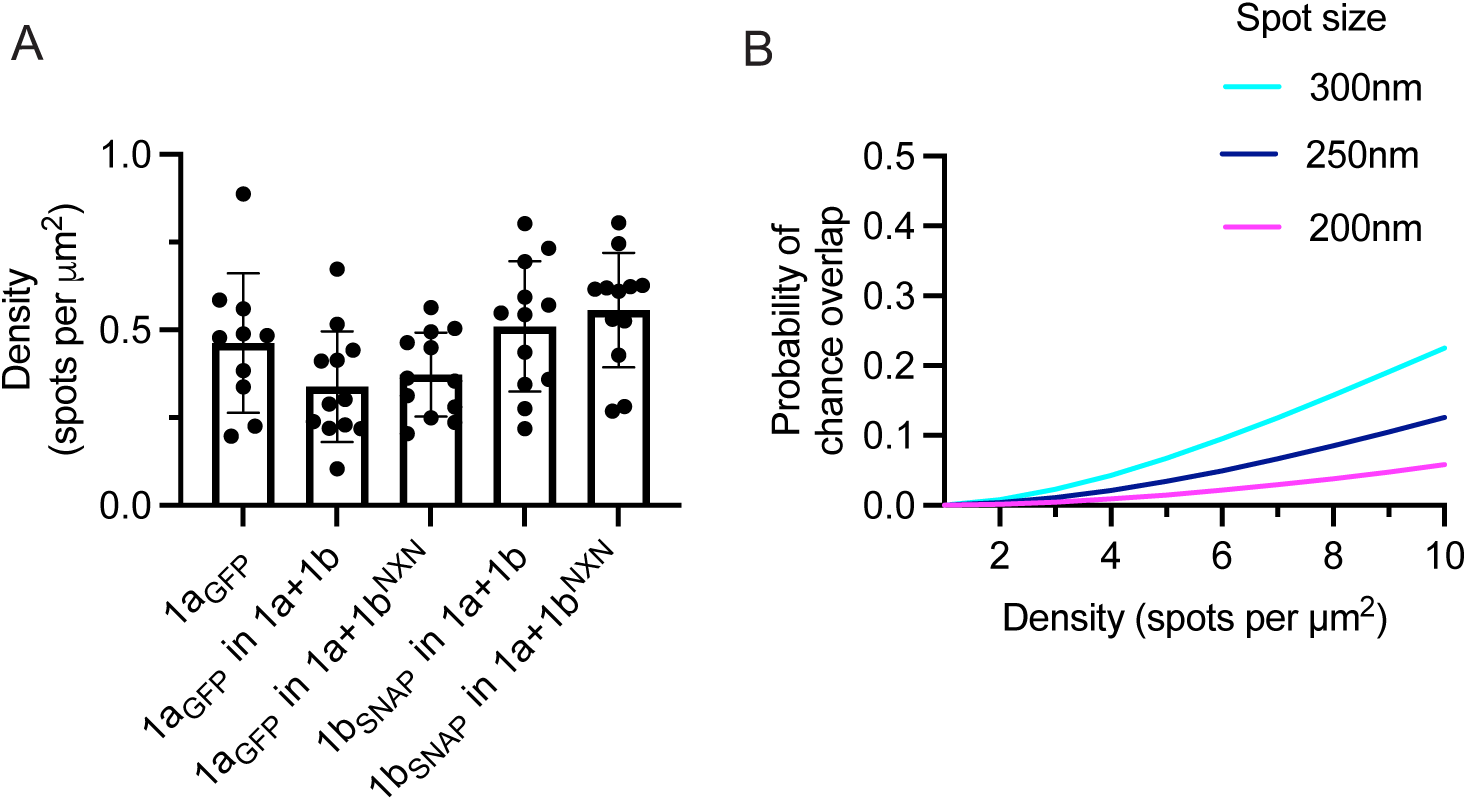
Density and probability of overlap by random chance. (*A*) The density of spots in the imaged cells used for stoichiometry analysis. (*B*) Probability of random chance of overlapping as a function of density, i.e., number of spots per μm^2^ calculated for different spot sizes; For density ∼0.5 μm^2^ the chance of overlap by random chance is almost zero.

